# Automappa: An interactive interface for metagenome-derived genome bins

**DOI:** 10.1101/2023.08.25.554826

**Authors:** Evan R. Rees, Samantha C. Waterworth, Shaurya C. Chanana, Jason C. Kwan

## Abstract

**Background:** Studies attempting to observe microbes commonly considered uncultivable under standard laboratory conditions are turning to so-called “deep” environmental sequencing approaches whereby they may access these unculturable organisms’ genomes via *in silico* approaches. A typical workflow involves metagenome assembly, annotation, and binning for reconstruction of each respective organism’s genome (or metagenome-assembled genome, MAG). Many automated genome binning approaches have been developed and have displayed a wide range of variation in performance. Therefore, refinement methods have been developed in order to aid manual curation following the automated genome binning process. Current manual curation tools were developed with a focus towards teaching metagenomics concepts and may fail when handling complex datasets containing many microbes. Automappa was developed with a focus on overlaying a variety of annotations such as taxonomy, coverage and marker-gene prevalence while maintaining an implementation that may scale to the complexity of environmental samples.

**Results:** We present Automappa, a companion tool and interactive interface for exploration and refinement of Autometa taxon and genome binning results from metagenomes. Selections provide real-time updates of MAG metrics to aid manual curation. Furthermore, researchers may detect unbinned MAGs as well as manually improve their draft-quality MAGs with contigs that closely match the MAG’s genome characteristics. Automappa’s utility has previously been demonstrated on host-associated, marine and terrestrial systems with a total of 242 curated MAGs across fourteen published metagenomes. Of these refined MAGs, the number of high-quality and medium-quality bins increased, consequently lowering the number of low-quality bins and decreasing the amount of data discarded from downstream analyses. The recovery of higher quality MAGs improved the confidence in results and strengthened the resultant conclusions of these respective studies. Automappa consists of three tabs, one for uploading a user’s metagenome data, another for exploration and refinement and the last for providing an overall summary of the refined MAG results.

**Conclusions:** Automappa is an open source software package that allows researchers to easily assess and refine undetected or draft-quality MAGs from their respective metagenomes. It is freely available under the GPLv3 license at https://github.com/WiscEvan/Automappa and through Figshare (doi: 10.6084/m9.figshare.22593235).

## Background

Innovations in next-generation sequencing methods have democratized studies reliant on analysis and exploitation of complex microbial communities. As sequencing continues to become more cost-effective, an even greater influx of researchers will employ these technologies. Un-targeted environmental studies have vastly expanded the diversity represented in the Tree of Life and consequently revealed many limitations in microbial cultivation [1–5]. However, whole genomes can be obtained through shotgun metagenomics (random direct environmental DNA sequencing), and this may lead researchers to insights regarding primary and secondary metabolism, ultimately aiding culturing efforts [6]. Such genomes obtained through metagenomics are termed metagenome-assembled genomes, or MAGs, and their quality is estimated through the presence of specific genes universally conserved in microbes [7]. Researchers may access *who* is in a particular sample as well as each species’ genetic capabilities at the same time, affording meaningful information across a wide range of industries. For example, in detection of agricultural pathogens, human-health monitoring through city waste systems and drug discovery efforts in de-orphaning therapeutically-relevant biosynthetic gene clusters (BGCs). Metagenomics is particularly well-suited for host-associated systems where culturing efforts have been deemed intractable and have allowed researchers to unravel previously misunderstood symbiotic relationships [8, 9]. The democratization of next-gen sequencing has generated “big data” and is accompanied with the development of a panoply of automated approaches.

The general computational workflow for genome-resolved metagenomics involves assembly of DNA sequencing data, various sequence annotation methods, binning of these annotated sequences into genome groups (MAGs) and finally characterization of the binned MAGs. In this context, a flawless automated approach should recover fully-assembled and classified genomes from the metagenome, eliminating any requirements of expert analysis or post-processing. However, genome binning of complex samples has remained recalcitrant with current state-of-the-art methods, limited in their ability to recover more than half of the members in both marine samples and communities with high amounts of strain-level overlap [10]. Achieving high-quality assemblies may require more sequencing, often including long-reads, which may be cost prohibitive to research groups. Furthermore, current long read technologies require sequencing coverage as well as DNA quality and quantity that may be unobtainable due to the scarcity of the sample. Consequently, one approach to improve the total recovery of MAGs has been to determine the best representative MAGs by consensus of various binners’ predictions with programs called “binning refiners” [11–13]. These binning refiners have been developed to circumvent the individual limitations across different genome binning methods. Binning refinement in this manner is constrained by the sensitivity of the aggregate binners’ predictions and consequently will fail to recover any members not initially recovered. Due to these limitations, many environmental metagenomics analyses often require expert curation.

Given current limitations, tools enabling manual analysis by researchers become indispensable. To this end, software is available to guide refinement of MAGs from metagenomic samples following (or in place of) an automated approach. There currently exist such tools as VizBin, ICoVeR and uBin that allow visualization of various metagenome annotations to provide a means to manually deconvolute microbes from their community [14–16]. VizBin provides a 2D scatterplot of metagenome sequences visualized as points corresponding to their dimension-reduced tetra-nucleotide frequencies [15] and the interface allows a user to select groupings of sequences to correspond to putative MAGs. ICoVeR allows verification and refinement with a set of interactive plots displaying annotations of sequence abundance, nucleotide composition and predictions from other binners. Additionally, ICoVeR provides automated refinement algorithms to further guide the MAG refinement process [16]. Similar to ICoVer, uBin supplies the end-user with a multitude of charts and graphs to curate a genome from a metagenome, although the intended purpose for uBin is to aid teaching metagenomics to students by familiarizing them with common metagenomic annotations [14]. However, all these software packages are limited to datasets less than approximately 90,000 contigs, and are limited in their user-customization options.

To overcome these problems and to aid our own binning efforts, we have developed Automappa. This is a companion tool to its automated binner, Autometa, which was constructed to address recovery of host-associated microbes [17]. Autometa is an automated workflow which aims to scale to the most complex communities that have been assembled. Therefore, Automappa was implemented to handle manual curation of MAGs at the scale of these complex datasets. Similarly, to address the aforementioned shortcomings of automated binning approaches, Automappa gives the user the ability to detect these divergent community members using a variety of visualizations. Previous binning refiners depict metagenomic data using GC content, contig coverage, taxon-binning annotations as well as nucleotide composition differences through the use of the dimension reduction technique, Barnes-Hut *t*-distributed stochastic neighbor embedding (tSNE). Automappa extends these metagenomic representations with the inclusion of an assortment of recently developed dimension reduction methods such as Uniform Manifold Approximation (UMAP) [18], the density-preserving versioof Pharmacy, University of Wisconsin - Madisonn DensMAP [19] and TriMap [20], a reduction method that uses triplet constraints to form the low-dimensional representation. Furthermore, metadata such as marker gene and taxonomy information are layered on top of the various embeddings to assist curation across all recovered MAGs.

## Implementation

Automappa was developed as a companion tool to the binning software, Autometa [17]. It was designed to visualize, verify and refine genome binning results to aid curation of high-quality MAGs. It is composed of interactive and inter-connected tables and figures that support selection with real-time MAG quality updates. The Automappa web user interface is implemented in Python and packaged with a docker-compose configuration file for ease of setup. Automappa was implemented to scale to metagenomics-sized data by connecting a variety of production-quality services, and it may be deployed on a lab server or an individual’s personal computer. Multiple services communicate with the Automappa server, allowing distribution and asynchronous execution of long-running metagenome annotation tasks. A Celery task-queue distributes tasks with RabbitMQ and Redis to manage the user’s input metagenome annotations [21–23]. These annotation tasks include a *k*-mer frequency analysis pipeline and retrieval of various MAG summary statistics. The back-end is managed by a Flask server, Redis, RabbitMQ and PostgreSQL while the front-end uses the Dash framework provided by Plotly [24–26]. Automappa is distributed with a Dockerfile for easy construction of an isolated server environment for use with various compute systems. The interface consists of three tabs. The first is the home page, where the user may upload results from Autometa’s automated binning workflow (Fig. 1). The second is the MAG refinement page, where the user can explore binning results and curate MAGs to improve overall MAG quality (Fig. 2). The third tab is a MAG summary page, where the user can access an overview of binning results as well as get an overview of any specific MAG of interest (Fig. 3).

**Figure 1.**
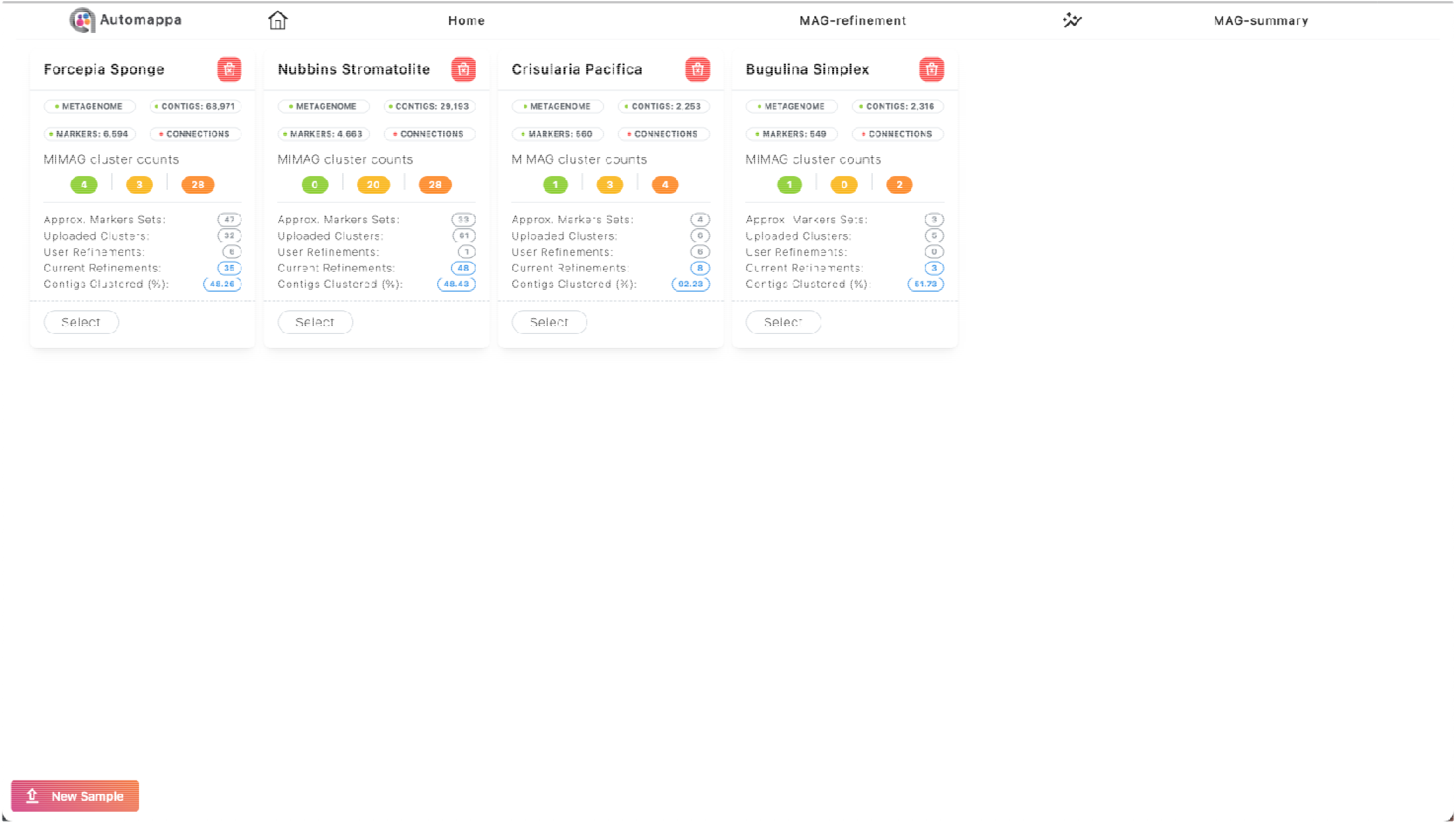
Home tab with four example datasets uploaded. Each sample card displays information with respect to the dataset, such as number of sequences, number of markers, number of initial MAGs uploaded and the total MAGs currently available. A breakdown of MAG counts corresponding to MIMAG quality are also determined for the currently available MAGs in the corresponding sample. These sample cards serve as an overview to the user for surveying their available datasets and refinement progress.

**Figure 2.**
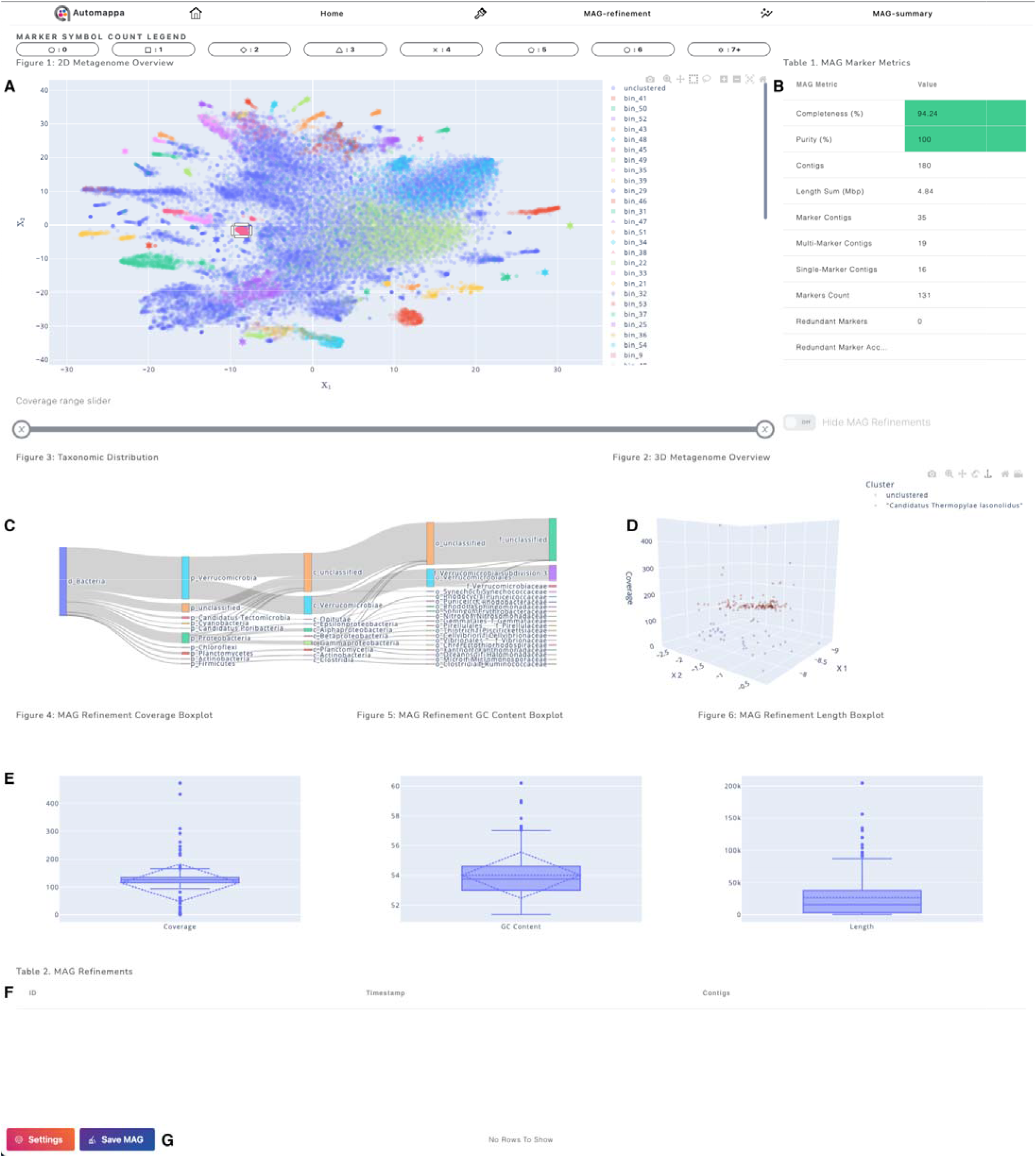
MAG refinement tab with uploaded *Forcepia* sponge data [27] displaying X_1_ and X_2_ embedded *k*-mer axes selected. (A) Points colored blue in the top left are unclustered contigs while the other colors represent specific MAG assignments. (B) The computed MAG marker metrics of the user-selected points from the 2D scatterplot. (C) The taxonomy designations from superkingdom to species canonical rank are directly below the top left scatterplot. By default this reflects all points except when a subset is selected. (D) To the right of the taxon binning annotations is a 3D scatterplot with the same x and y axes as the 2D scatterplot and coverage for the z axis. The 3D scatterplot automatically updates to show the selected points only. (E) Boxplots depicting coverage, GC content and length distributions are available, after selection (similar to the MAG summary in Figure 3, below). (F) MAG refinements table containing information of user refinements. (G) Two buttons affixed for changing figure settings (for example, mapping of axes to different variables), deleting, downloading and saving user’s MAG refinements. This view is accessed directly after navigation to the *MAG Refinement* tab, so all contigs are represented across the figures as well as within the MAG Marker Metrics table. Selection of a subset of contigs in the 2D plot results in other figures and tables updating to show information for the selected contigs only as is shown for the group of points encompassed by the rectangle in A.

**Figure 3.**
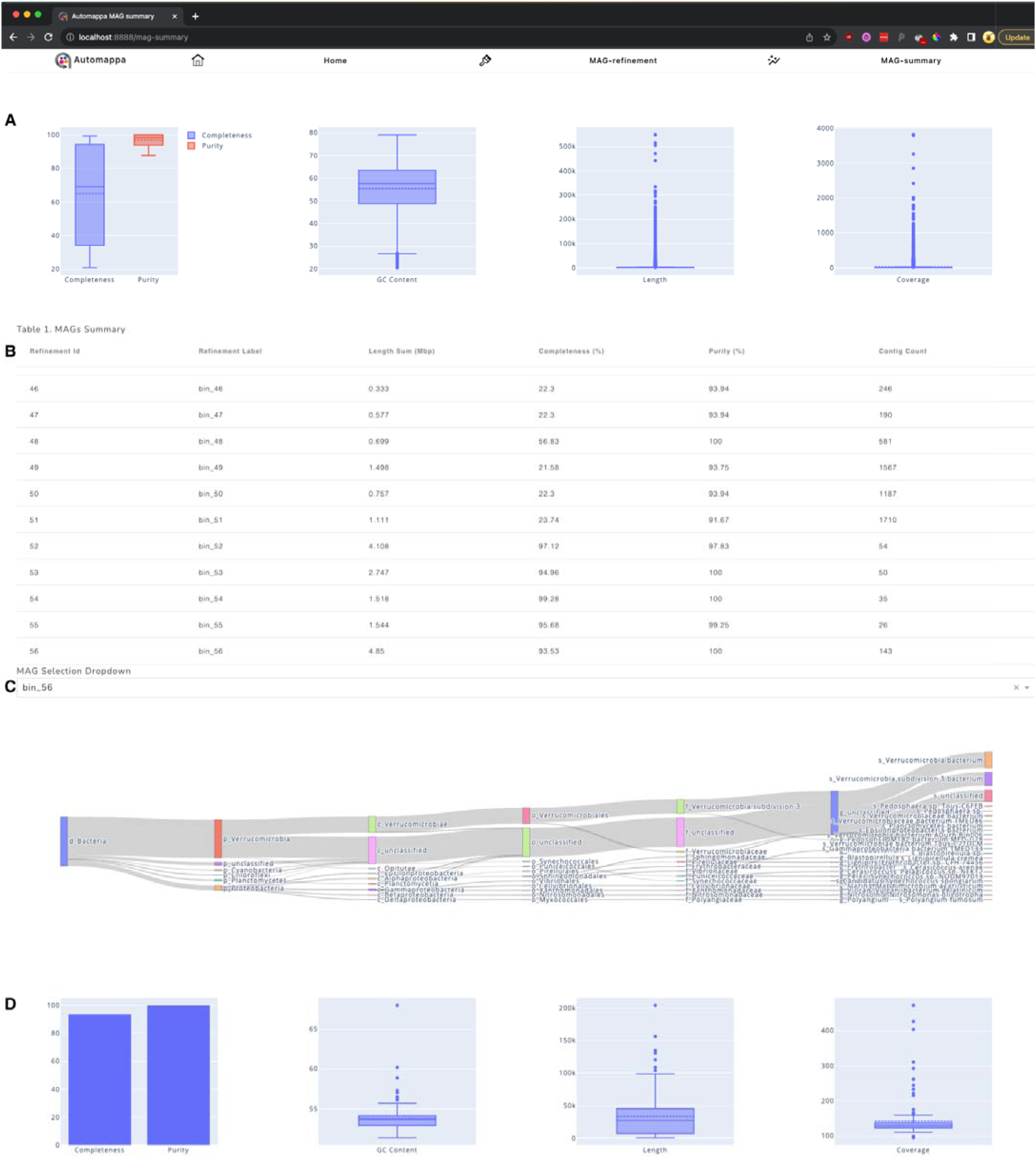
MAG summary overview. (A) Box plots showing completeness, purity, GC content, length and coverage of all MAGs both imported and refined. (B) Table of MAG characteristics, including total length, estimated completeness and purity. (C) A dropdown for MAG selection which will populate the Sankey diagram showing the distribution of taxonomic classifications at each rank for the selected MAG. (D) Characteristics of the selected MAG represented through bar and box plots.

The home tab allows the user to upload their dataset and displays an assortment of sample cards indicating various statistics determined for each of the user’s samples. The cards are persistent between runs, so data does not have to be uploaded each time Automappa is started. Three files are required for upload: (1) the main binning annotation file written by the autometa-binning command using taxonomy information, (2) the markers file written by the autometa-markers command and (3) the corresponding fasta formatted metagenome assembly. Upon upload, annotations are ingested into a PostgreSQL database and then their files are removed. After this, the user may select these data by selecting the corresponding sample card for use during the MAG refinement process. Subsequent navigation throughout Automappa will use the data selected at this step.

The main view of the MAG refinement tab is composed of multiple connected and interactive figures (Fig. 2): (A) A 2D scatterplot of contigs where the user may specify each axis reflecting a variety of contig annotations. The default axes are contig annotations of GC content and coverage. Additional annotations of embedded *k*-mer coordinates are computed upon dataset selection. These coordinates are determined by a *k*-mer frequency analysis workflow. Briefly, a window of size *k*, is used (typically 4 or 5) to construct all possible combinations of nucleotides at the given *k*. These *k*-mers are searched and counted for each contig then all counts are normalized by a center-log ratio transformation. Finally the normalized *k*-mer frequencies are processed by a dimension reduction technique, also known as embedding, to project the high-dimensional *k*-mer frequencies down to two dimensions, shown as “X_1_” and “X_2_” in Automappa’s settings pane. In this embedded space the relation of one contig to another represents the similarity or difference by nucleotide composition. The user can change the scatterplot’s axes within the settings menu between either GC content vs. coverage or X_1_ vs. X_2_, and can specify that points be colored by cluster or specific taxonomic rank. Contig marker annotations are overlaid in this scatterplot based on their marker count to aid the user in the MAG refinement process. Users are able to select groups of contigs using either rectangular or lasso selection tools, and in addition there is a slider underneath the main 2D plot that hides points outside of the specified coverage range, allowing for selection within that range only. (B) A datatable summarizing the characteristics of the selected contig group, including completeness and purity inferred through single-copy marker genes. (C) A Sankey diagram to visualize each selected contig’s taxon distribution across each canonical rank from superkingdom to genus. (D) A 3D scatterplot displaying the selected coordinates from the 2D scatterplot (utilizing the same x and y axes) with an additional axis of contig coverage, length or GC content. Coverage in this context is especially useful for resolving samples with strain-overlap or for observing possible incidences of horizontal gene transfer. (E) Box plots to assess the distribution of MAG characteristics such as GC content, coverage and length in the selected points. Distributions of GC content and coverage help the end-user to determine whether the possible MAG refinement is feasible, while the MAG refinement assembly quality can be inferred by looking at the distribution of fragment lengths. Most useful regarding the MAG refinement page is the interconnectedness of these figures and the associated tables. Contigs may be selected in the 2D scatterplot causing the other figures to be updated based on the selection. Furthermore, MAG completeness and purity measures are updated in real-time with additional single-copy marker metrics displayed alongside the figures. This allows the user to manually curate MAGs with selections guided by repeated measures of increasing completeness and decreasing contamination (or increasing purity). When the user has finalized their curation, they may save their manual refinements which can be exported when they are finished with the refinement process. To aid the user during the refinement process, a table at the bottom of the page is updated with the user’s MAG refinements. To avoid accidental duplication of a contig’s membership into multiple MAGs (and potentially erroneous interpretation of results), a contig is assigned to its latest MAG classification and removed from its previous MAG assignment. Contigs that have already been placed in a manually curated MAG may be hidden by a toggle located at top of the page. When the user has resolved all of their MAGs and has deemed the refinement process as complete, they may navigate to the MAG summary page for a general overview.

The MAG summary page (Figure 3) provides aggregated statistics of the recovered MAGs to easily assess binning performance. The page is composed of various figures and a MAG summary table. Upon navigation to the MAG Summary tab, four boxplots are generated providing an overview of binning results and metagenome information. The first box plot displays the distribution of completeness and purity estimates across all of the recovered MAGs. The other three boxplots are updated with metagenome distributions of GC content, length and coverage. Underneath these boxplots is a MAG summary table which provides aggregated MAG-specific statistics available to the end-user for review and scrutiny. These statistics allow the user to critique a MAG’s fragmentation or assembly quality (N50, N90, N10, size, contig count), marker gene estimates and length-weighted distributions of coverage and GC content.

The user may inspect an individual MAG by use of the MAG selection dropdown beneath the MAG summary table. Upon MAG selection, a Sankey diagram of the respective MAG’s contig taxonomy distribution across canonical ranks is generated neighboring three additional box plots to assess the MAG-specific distribution of GC content, length and coverage with their corresponding quartiles. A bar plot of the MAG’s completeness and purity is also created adjacent to the aforementioned boxplots, as shown in figure 3D. When a user has completed their MAG refinements, they can be downloaded through the Settings menu under the MAG refinement page.

## Results and discussion

Several automated genome binning algorithms have been developed, each striving to produce the highest quality MAGs possible. However, these tools fail individually and in aggregate to recover complete and pure metagenomic communities [10]. Some of these tools may exhibit a black-box effect, where the user may not fully understand how their final MAGs were binned due to the obscurity of the genome binning methods. For example, a *k*-mer frequency analysis is a measure of nucleotide composition which first requires counting *k*-mers of length *k*, leading to a high-dimensional distribution of counts which are then normalized before being reduced to two dimensions via a non-linear dimensionality reduction technique, such as tSNE. Consequently, the dimension-reduced representation of the original similarities and differences within the high-dimensional *k-* mer counts becomes obscured. Additionally, recursive binning predictions may further obfuscate a method’s decision-making, limiting the user’s ability to discern the properties of why sequences were clustered as they were. Manual curation of metagenomic bins gives researchers control over their final resolved bins, for example through inclusion of unclustered contigs, splitting of MAGs by abundance or coverage, or combination of incomplete MAGs to produce higher quality bins.

We suggest users take the following deliberate approach to manually curating their clusters: Within the MAG Refinement tab, moving down the cluster legend of the 2D scatterplot in the upper left, one can iteratively manually curate each cluster by 1) examining the 3D plot to determine if coverage is approximately uniform across the cluster. If a cluster appears to have a stark difference in coverage (as seen in the 3D plot, previously mentioned), it is recommended that this bin be split into two or more clusters by utilizing the coverage slider underneath the 2D scatterplot, which may result in improved gene marker purity. 2) If coverage appears uniform, the user can proceed to assess whether combining the cluster with a neighboring MAG or surrounding unclustered contigs improves completeness with minimal or no change to the purity levels. Alongside coverage and completeness, the user may also inspect uniformity between the contig selection’s taxonomic distributions. This is done by iteratively selecting different contig groupings. For example, inclusion or exclusion of marker-containing contigs (indicated in Automappa with distinct shapes corresponding to their respective marker counts outlined by the *Marker Symbol Count Legend* above the 2D scatterplot in Figure 2A) will alter the MAG quality completeness and purity metrics as well as potential change the taxonomic composition of the selection if there are contaminants. As Autometa errs on the side recovering incomplete clusters of high purity, improvement of bin quality can often be achieved by the addition of unbinned contigs or the combination of two or more clusters. This process, at the expense of introducing limited contamination (below 3% contamination per bin is advised), can achieve large increases in MAG completion. Toggling between coloring contigs by genus or species may also be useful in determining which contigs to include in a given cluster. When the user decides their refinements represent the highest quality MAGs possible, they can save the selection and move on to the next MAG refinement.

With this approach, Automappa has been used in our group to manually curate 242 MAGs from fourteen metagenomic samples across a diverse range of environments, including stromatolites, marine sponges and beetles which have been successfully published in scientific literature [8, 27–29]. Moreover, this tool has been used to produce an additional 376 final clusters from 13 more unpublished metagenomes derived from a similar diverse range of environments. Of these metagenomes were sponge (Phylum *Porifera)* microbiomes within which a genome representing a novel *Tethybacterales* family was recovered as well as eleven other genomes in the *Tethybacterales* order [29]. The refined MAGs comprising these broadly-distributed *Tethybacterales* families were compared to discern any potential functional differences. Automappa assisted in gaining these insights by enabling expert curation of metabolic pathways respective to each of the *Tethybacterales* MAGs. Similarly, microbiomes from geographically distinct stromatolite formations were curated using Automappa allowing comparison of the conserved MAGs’ functional gene sets recovered [28]. Automappa was also utilized to better understand a terrestrial host-associated metagenome. MAGs were curated from the eggs of the *Lagria villosa* beetle, leading to the surprising discovery of a reduced genome symbiont, *Burkholderia* sp. LvStB, as the exclusive producer of the antifungal compound, lagriamide [8]. Furthermore, multiple horizontally transferred genes were found within the genome of strain LvStB, shedding light on the initial stages of the evolutionary process of genome reduction. More recently a sponge (Phylum *Forcepia*) microbiome was analyzed and with Automappa assisted in identifying *“Candidatus* Thermopylae lasonolidus”, a candidate producer of lasonolide A and helped to suggest the potentially horizontally transferred lasonolide A biosynthetic gene cluster using pentanucleotide frequency and GC content, which are readily available for inspection in the MAG refinement view [27].

## Future development

All current automated genome binning approaches fail to recover greater than fifty percent of the genomes from any of the most recent CAMI2 challenge datasets [10]. These datasets consisted of a community with high amounts of strain overlap and a marine metagenome, containing many less represented taxa in the NCBI databases. Future methods will need to address the complexities of separating closely-related genomes as well as for handling highly-divergent, erroneous database matches. Not only must binners account for these challenges, but they must also be capable of scaling to highly populous communities, such as collections of soil samples. Forthcoming developments within the Autometa workflow will attempt to address these intricacies. Automappa will pursue follow-up implementations to continue to aid the end-user in exploration and refinement of their sample, regardless of environment. Some of the planned future developments include integration of pre and post-processing workflows such as genome binning with Autometa, taxon binning with the Genome Taxonomy Database Toolkit (GTDB-Tk), lineage-resolved MAG quality analysis using CheckM [30, 31] and addition of MAG-specific KEGG annotations using tools such as kofamscan [32] or METABOLIC [33]. Outside of incorporation of commonly used comparative genomics tools, additional interactive figures and features embedded within these interactive features are outlined. For example, with datasets containing more than 100,000 sequences, overplotting may become problematic. To reduce overplotting, putative MAGs may be consolidated by visualizing their geometric median thereby dually increasing app performance and better differentiating consolidated MAGs with the surrounding unclustered contigs. Supplementary to selections whereby MAG metrics are iteratively improved, an interactive plot of contigs’ assembly graph connections may be added allowing the user to easily apprehend the respective MAGs inter and intra-MAG connections. This would help the end-user to determine possible horizontal gene transfer events and help to resolve contigs with highly different coverage profiles. Automappa is open-source and has undergone multiple iterations with features requested by its users. The core Automappa framework allows extension of additional features that are requested as Automappa is continually employed on a variety of datasets from distinct environments.

## Conclusions

We present Automappa, an interactive interface for exploration and refinement of metagenomes. The application is capable of recovering and improving undetected and draft-quality MAGs. Automappa is open-source and freely available to be deployed on a lab server or personal machine. It has been designed to allow integration of post-processing steps to further facilitate the end-user’s interpretations and conclusions for their respective comparative genomic studies. Automappa was conceived out of necessity to enable lab-members to carry out metagenomic analyses where current fully-automated state-of-the-art metagenomics workflows fail to resolve members from complex microbial communities. We previously demonstrated Automappa’s utility by manual curation and refinement of MAGs across marine, terrestrial and host-associated metagenomes. These refinements lead to improved downstream functional and evolutionary analyses enriching the insights and conclusions respective to each study. We also suggest future improvements to the application as well as integrations of post-processing comparative genomic tools to further enable experts during their metagenomics workflow.

## Availability and Requirements

Project name: Automappa

Project home page: https://github.com/WiscEvan/Automappa

Operating system(s): Platform independent

Programming language: Python

Other requirements: Python >=3.9

License: AGPL-3.0 License

Any restrictions to use by non-academics: None

## List of abbreviations

MAG: Metagenome-Assembled Genome
CAMI: Critical Assessment of Metagenome Information
GTDB-Tk: Genome Taxonomy Database Toolkit

## Competing interests

The Kwan lab is planning to offer their metagenomic binning pipeline Autometa on the paid bioinformatics and computational platform BatchX in addition to distributing it through open source channels.

## Funding

This work was supported by the U.S. National Science Foundation (DBI-1845890) and the Gordon and Betty Moore Foundation (Grant number 6920).

## Availability of data and materials

All data and materials are available as supplementary files or can be downloaded from the software website https://github.com/WiscEvan/Automappa. Automappa is released under GPLv3 license.

## Consent for Publication

Not applicable

## Ethics approval and consent to participate

Not applicable

## Authors’ contributions

ER, SW and SC designed the application. ER and SC created the application. ER and SW wrote the manuscript. JK coordinated the study and helped to draft the manuscript. All authors read and approved the final manuscript.

## Notes

https://github.com/WiscEvan/Automappa

https://figshare.com/articles/software/Automappa/22593235

